# Skill Acquisition is Enhanced by Reducing Trial-To-Trial Repetition

**DOI:** 10.1101/866046

**Authors:** Lore WE Vleugels, Stephan P Swinnen, Robert M Hardwick

## Abstract

Developing approaches to improve motor skill learning is of considerable interest across multiple disciplines. Previous research has typically shown that repeating the same action on consecutive trials enhances short-term performance, but has detrimental effects on longer term skill acquisition. However, most prior research has contrasted the effects of repetition only at the block level; here we examined the effects of repeating individual trials embedded in a larger randomized block a feature that is often overlooked when generating random trial orders in learning tasks. With four days of practice, a “Minimal Repeats Group”, who rarely experienced repeating stimuli on consecutive trials during training improved to a greater extent than a “Frequent Repeats Group”, who were frequently presented with repeating stimuli on consecutive trials during training. Our results extend the previous finding of the beneficial effects of random as compared to blocked practice on performance, showing that reduced trial-to-trial repetition during training is favorable with regards to skill learning. This research highlights that limiting the number of repeats on consecutive trials is a simple behavioral manipulation that can enhance the process of skill learning. Data/analysis code and supplementary materials available at https://osf.io/p3278/

**NEW & NOTEWORTHY:** Numerous studies have shown that performing different sub-tasks across consecutive blocks of trials enhances learning. Here we examined whether the same effect would occur on a trial-to-trial level. Our Minimal Repeats Group, who primarily responded to different stimuli on consecutive trials, learned more than our Frequent Repeats Group, who frequently responded to the same stimulus on consecutive trials. This shows that minimizing trial-to-trial repetition is a simple and easily applicable manipulation that can enhance learning.

## INTRODUCTION

New skills require a considerable amount of practice to learn and master (Ericsson et al. 1993; Magill and Anderson 2013). Many studies have investigated different methods to improve the rate at which motor skills are learned, including approaches such as manipulating knowledge of results and other types of feedback (Debaere et al. 2003; Lauber and Keller 2014; Rhoads et al. 2014; Ronsse et al. 2010; Sharma et al. 2016; Swinnen et al. 1990; Wulf et al. 2010), manipulating explicit awareness (Kal et al. 2018; Robertson 2007), providing reward or punishment (Abe et al. 2011; Chen et al. 2018; Galea et al. 2011; Manohar et al. 2015; Wächter et al. 2009), and applying non-invasive brain stimulation (Galea et al. 2011; Hardwick and Celnik 2014; Reis et al. 2009; but see also López-Alonso et al. 2015, 2018; Vallence et al. 2013). A growing body of research has focused on the consequences of repetition on both the execution and learning of motor tasks (Fecteau and Munoz 2003; Gupta and Cohen 2002). Research on effects such as “repetition priming” (Gupta and Cohen 2002), “facilitation of return” (Tanaka and Shimojo 1996), and “priming of pop-out” (Fecteau 2007; Maljkovic and Nakayama 1994) all refer to implicit short-term memory phenomena which facilitate the processing of a stimulus as a result of exposure to this same stimulus in the preceding trial (Schwartz and Hashtroudi 1991). Such priming effects are linked with neuroscientific research on repetition suppression the finding that repeating the same stimulus or action leads to a reduction in trial-to-trial changes in neural activity (Desimone 1996; Hamilton and Grafton 2009; Henson and Rugg 2003; Wiggs and Martin 1998; Wymbs and Grafton 2015), as the circuits involved in producing the specific response to that stimulus or action are already primed for action.

Several previous studies have investigated the short-term consequences of repeating the same action from trial-to-trial. This has been shown across a diverse range of paradigms and measures of performance; reaction times are shorter on repeated trials when the time-lag between the stimulus presentation and response was short (Bertelson 1961), and when repeating stimulus features such as color or position (Maljkovic and Nakayama 1994, 1996; Tanaka and Shimojo 1996), hand reach trajectories are more curved when a participant had to avoid an obstacle in the previous trial (Jax and Rosenbaum 2007), and smooth saccadic eye movements are dependent on previous target motions (Kowler et al. 1984). Similar trial-to-trial priming effects are present when we observe the actions of another person prior to performing the same action ourselves (Edwards et al. 2003; Griffiths and Tipper 2009; Hardwick and Edwards 2011). Taken together, repetition on a trial-to-trial level has shown to modulate and enhance performance in the short-term.

The effect of repeating the same action has also been examined in learning. This is typically achieved by dividing the task to be learned into sub-tasks and manipulating the order in which these sub-tasks are practiced. For example, performing a jump serve in volleyball (i.e. task) can be practiced by executing the throw of the ball, the timing of the jump you will make, and the actual act of striking the ball (i.e. sub-tasks). Research has shown that “blocked practice” (practicing by repeating the same sub-task across consecutive blocks) leads to better performance during the acquisition of motor skills, whereas “random practice” (practicing different sub-tasks across consecutive blocks) leads to better retention (Battig 1979; Cross et al. 2007; Goode and Magill 1986; Pauwels et al. 2014; Smith and Davies 1995; for a review see Magill and Hall 1990). It is generally agreed that this "contextual interference" effect occurs because random practice is more challenging than blocked practice (Pauwels et al. 2014, 2018). Though the exact mechanism of action remains under debate (c.f. Coxon et al. 2012; Lee et al. 1985; Lee and Magill 1983; Shea and Zimny 1983; Shea and Morgan 1979), the beneficial effects of inducing contextual interference through random practice have been widely examined and replicated (Brady 1998; Magill and Hall 1990; Merbah and Meulemans 2011).

While numerous studies have investigated the effects of repetition on learning at the block level, this effect has received little examination at the level of individual trials. For example, Kaipa et al. (2016) investigated the effect of repetition of phrases in a speech learning task, but used only two phrases. Though recent work has generally shown beneficial effects of trial-to-trial repetition (Ariani et al. 2019; Mawase et al. 2018), these studies used single group, within-subject approaches; as such, they were not designed to examine whether varying the amount of repetition a participant completes during training affects their overall learning. Notably, many training protocols involve presenting different trial types in a (pseudo)random order without controlling for the amount of trial-to-trial repetition that occurs, despite the effects of such repetition being largely unexplored. Therefore, manipulating the amount of trial-to-trial repetition that occurs when learning a task has considerable potential as an easily implemented, purely behavioral approach to augment the learning process.

The present study investigated the effects of switching between and repeating the same response as participants learned an arbitrary visuomotor association (Hardwick et al. 2019). As repeating the same action facilitates performance, we hypothesized that training that included consecutive repetition of the action would represent a less challenging practice condition, effectively inducing a lower quality of training. We tested this hypothesis by comparing the results of two groups of participants; a Minimal Repeats Group that experienced different stimuli on the majority of consecutive trials, and a Frequent Repeats Group that experienced relatively more trial-to-trial repetition on consecutive trials. As the Minimal Repeats Group trained under a more challenging context, we predicted that they would show a greater improvement with training than the Frequent Repeats Group.

## MATERIALS & METHODS

### Participants

A total of 39 young and healthy participants were recruited for the study. Overall, data from eight participants were excluded from the study. Three participants were excluded as they did not properly follow the instructions (these participants did not consistently make responses at the same time as the fourth tone during forced response conditions; see Materials & Methods). A further five participants were excluded due to technical errors in data acquisition. This left a total sample of 31 participants (mean age = 21.7, age range = 18.4–25.3). Subjects were randomly assigned to one of two groups. The Minimal Repeats Group (n = 15, male = 5, female = 10, mean age = 21.8, left handed = 3), who trained with minimal repetition of consecutive trials (≈1% of trials), and the Frequent Repeats Group (n = 16, male = 5, female = 11, mean age = 21.6, left handed = 3), who were presented with repeated stimuli on ≈21% of consecutive trials during training (Figure 1b). Participants were not aware of the division of the groups. The two groups did not differ significantly with regards to handedness (t_29_=0.03, p=0.933), gender (t_29_=0.06, p=0.905), or age (t_29_=4.38, p=0.663). All participants gave written informed consent before the start of the experiment. The study was approved by the local ethical committee of the University of Leuven (KU Leuven), Belgium. Participants received financial compensation (€20) for their participation.

**Figure 1.**
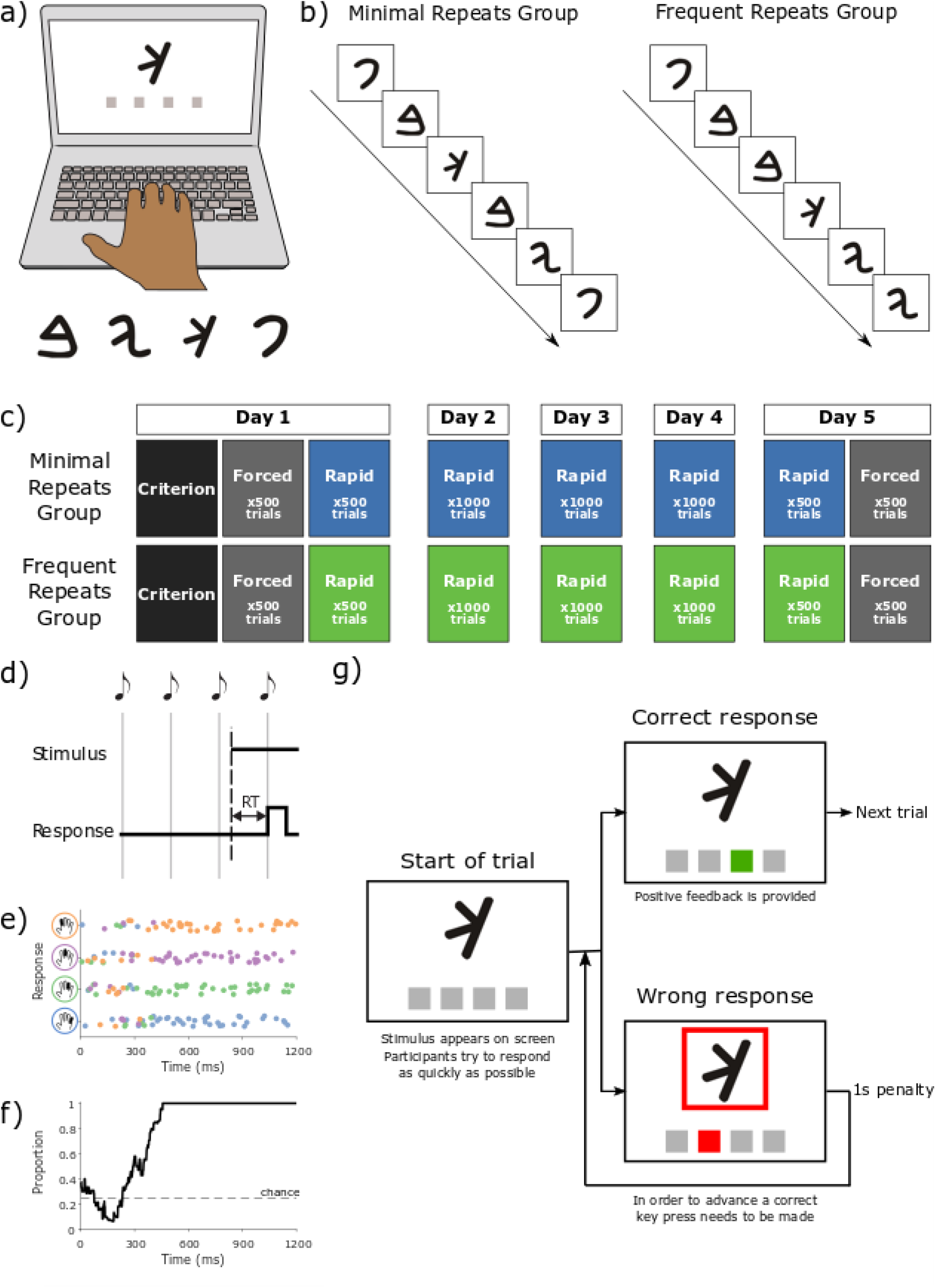
Task setup and procedure. a) <u>Task setup</u>. One of four stimuli (lower) was presented on screen (upper). Participants had to respond by pressing a corresponding key with a certain finger. b) <u>Groups</u>. Rapid response training blocks differed between groups. The Minimal Repeats Group rarely (~1% chance) experienced repeating stimuli on consecutive trials, whereas the Frequent Repeats Group were often (~20% chance) presented with repeating stimuli on consecutive trials. c) <u>Task procedure</u>. On day 1, participants came to the lab and were familiarized with the task (not presented in figure, see Materials & Methods - General Procedures for further details). Hereafter, in the criterion test block, they learned a fixed mapping between stimuli and keys that was used throughout the rest of the experiment. They then completed forced and rapid response trials. Note that only the rapid response training trials differed between group, as described above. On days 2, 3, and 4, participants trained on the task at home using rapid response trials. On day 5, participants came back to the lab. They first were presented with rapid response trials, after which their final level of performance was measured during forced response trials. d) <u>Forced response paradigm</u>. Participants heard a sequence of four tones, each separated by 400ms, and were instructed to make a response synchronously with the fourth tone. A stimulus appeared at a random time between the first and fourth tone, effectively imposing a limited response time. e) <u>Forced response example data</u>. Data from an example participant on the familiarization task. Participants responded to pictures of hands in which one finger was highlighted by pressing the corresponding finger on the keyboard. Illustrations of these stimuli are presented on the y-axis. Each dot represents the participant’s response on a single trial (jitter added to y-axis to allow for visualizing multiple responses at the same time). Each color represents the finger the participant responded with. Note that when the reaction time was low, all colors are mixed, indicating that the participant did not have enough time to prepare the correct response. However, when the time allowed to prepare a response was greater, the participant was able to make a correct response, as indicated by the consistency of the colored dots. f) <u>Data from panel c converted into a speed-accuracy trade-off curve</u>. A sliding window (running average across a 100ms window) was used to visualize the accuracy for a given time. Note that at relatively short (less than approximately 300ms) latencies, responses were around chance level, meaning that participants did not have enough time to process the stimulus and hence pressed a random key. After approximately 300ms, the level of accuracy increased with time until reaching a plateau. Note the relationship between the switch from random to consistent selection of the appropriate response in panel c corresponds to the proportion of correct responses in panel d. g) <u>Rapid response paradigm</u>. A stimulus appeared on screen and participants had to respond as quickly as possible by pressing one of the four keys on their keyboard. Whenever a participant made a correct response they advanced to the next trial (300ms delay between trials). However, when a participant made an incorrect response, there was a 1000ms delay (“1s penalty”) before they could make another response. Participants had to provide the correct response in order to advance to the next trial.

### General Procedures

Participants performed a finger pressing task (Arbitrary Visuomotor Association Task; Hardwick et al. 2019) on their personal computer. They had to respond to the appearance of one out of four stimuli by pushing a specific key (F, T, Y, or J) on their computer keyboard with either the index, middle, ring, or little finger of their dominant hand (Figure 1a). Four different types of conditions were completed: familiarization, criterion test, rapid response, and forced response conditions. In all conditions, stimuli were presented in pseudorandomized sub-blocks of 20 trials, with each stimulus appearing five times within each sub-block. The familiarization, criterion test, and forced response conditions were the same for both groups. This was not the case in the rapid response training condition (see Figure 1b-c). In this condition, for the Minimal Repeats Group, a different stimulus was presented for each consecutive trial within these sub-blocks. For the Frequent Repeats Group, each stimulus repeated once on consecutive trials within each sub-block (i.e. 4/20 or 20% of trials presented repeat stimuli within each sub-block). There was a 25% chance of a repeat from one sub-block to another (i.e. one sub-block ending with the same stimulus that began a new sub-block). As each block comprised five sub-blocks, there were therefore four transitions between sub-blocks. Consequently, there were approximately 1 in 4 repetitions between the sub-blocks, meaning there was approximately 1 repeat between sub-blocks per overall block of 100 trials. This helped reduce the likelihood that participants would become aware of the experimental manipulation (i.e. if participants realized they could eliminate the previously pressed button from contention, their baseline level of accuracy would improve from 1/4 to 1/3).

#### Familiarization tasks

Familiarization trials provided participants with a chance to learn the requirements of the task. Here participants had to respond to pictures of hands, rather than arbitrary stimuli. In these pictures, one finger was highlighted, and participants had to respond with the corresponding finger. During these familiarization blocks, participants performed trials in both rapid response and forced response conditions, each of which are explained in more detail below.

#### Criterion test block

In this block, participants learned the correct mapping of the keys and stimuli through trial and error. A stimulus appeared on screen and participants had to learn which key it corresponded to. Stimuli were one of four different symbols, and each symbol corresponded to a different key. The mapping of the keys and stimuli was counterbalanced across participants. During this block, participants were instructed to focus on learning the correct mapping of the symbols and keys. Therefore, they were instructed that they could take as much time as they needed (i.e. there were no reaction time constraints). The block ended as soon as a participant had made 5 consecutive correct responses to each of the four stimuli (i.e. 20 trials in total). When the participant responded correctly to a stimulus they received positive audiovisual feedback (the on-screen box corresponding to the pressed key turned green, and a tone indicating the correct response played), the task continued to the next trial following a 300ms delay. Incorrect responses were met with negative audiovisual feedback (a red box was presented around the stimulus, the on-screen box corresponding to the pressed key turned red, and an unpleasant buzzer sound was played). Following an incorrect response, the participant had to wait 1000ms before their next response was registered.

#### Rapid response (training) blocks

During rapid response blocks (see Figure 1g), participants had to complete trials by responding to the stimulus as quickly as possible using the corresponding finger. The goal was to complete the block as quickly and as accurately as possible. As in the criterion assessment trials, whenever participants made a correct response (which was made clear by providing feedback, i.e. the on-screen box corresponding to the key that was pressed turned green, and a tone indicating the correct response played), they continued with the next trial after a 300ms delay. However, if a participant made a wrong response, they received audiovisual feedback (a red box appeared around the stimulus, the on-screen box corresponding to the key that was pressed turned red, and an unpleasant buzzer sound was played) and the participant had to wait 1000ms before another response could be made (referred to as a “one second penalty” when explaining the condition of the task to the participants). Participants were instructed to respond as quickly as possible, but to find a balance between speed and accuracy. At the end of each rapid response block an on-screen display showed the overall time required to complete the block, and compared this with their ‘personal best’ block completion time, encouraging participants to continually improve this personal best time.

#### Forced response (assessment) blocks

During forced response trials, participants heard a series of four equally spaced tones (400ms separation between each tone), and were instructed to time their response so that it would coincide with the fourth tone (Figure 1d). In each trial a stimulus appeared at a random time between the first and fourth tone. This allowed effective control over the amount of time participants had to process the stimulus and prepare their response. The earlier the stimulus appeared on screen, the more time the participants had to prepare their response. On the contrary, the later the stimulus appeared on screen, the less time the participants had to prepare their response, and the more difficult it was for them to make a correct response. Participants were instructed that their highest priority was to respond at the same time as the fourth tone, and that while they should aim to be accurate wherever possible, this should not come at the cost of missing the deadline; as such, their likelihood of success in trials in which the stimulus appeared at a time too late for them to accurately process was approximately 1 in 4 (Figure 1e,f). In contrast with the rapid response and criterion test conditions, no time penalties were enforced for providing incorrect responses (these penalties would encourage participants to ignore timing demands and instead focus on taking longer to provide an accurate response that avoided a penalty). Participants received on-screen feedback informing them that they had responded ‘too early’ or ‘too late’ if they responded more than 100ms before or after the fourth tone, respectively. For each trial in which a response was registered, the ‘preparation time’ for that trial was calculated as the time between the presentation of the stimulus and the time at which the participant responded. As such, trials in which participant responses were outside the ±100ms window were not discounted (as the actual time the participant used to prepare their response could still be accurately measured).

## PROTOCOL

### Day 1

Session 1 started with a brief introduction to the task. Participants first completed two familiarization conditions (2 blocks of reaction time based rapid response trials, followed by two blocks of forced response based trials, 100 trials per block, all to non-arbitrary hand stimuli). After the task familiarization trials, participants then completed a criterion test block in which they learned a fixed mapping between the experimental stimuli and specific keys. This mapping remained fixed for the rest of the experiment. Subsequently, participants executed 5×100 trial blocks of forced response trials, allowing us to collect a baseline speed-accuracy trade-off. Finally, participants trained on 5×100 trial blocks of rapid response trials.

### Days 2 – 4

Participants completed reaction time based rapid response training trials on the second, third, and fourth session. Each session consisted of 10 blocks, each comprising 100 rapid response trials to arbitrary stimuli.

### Day 5

The first five blocks were rapid response trials to arbitrary stimuli, allowing a measure of performance on the final day of training with this trial type. The final five blocks were forced response trials to arbitrary stimuli, collected in order to determine the effects of training on the participants’ speed-accuracy trade-off.

## DATA ANALYSIS

### Criterion Test Trials

During the criterion test block, two measures were extracted. Firstly, the number of trials participants needed to reach the criterion level (i.e. the number of trials a participant needed to make five consecutive correct responses to each of the four stimuli) was measured and compared between groups using an independent samples t-test. While participants were explicitly instructed that there were no reaction time constraints in this condition (i.e. they could take their time to complete the block and did not have to respond as fast as possible), we also conducted an analysis on their reaction times in order to fully quantify their performance. Reaction times for all trials in the criterion block were compared between groups using a linear mixed model with the form Reaction Time ~ Group+(1|Subject).

### Forced Response Trials

During the forced response trials, we determined ‘preparation time’ as the time between the presentation of the stimulus and the time of the participant’s response, and also recorded whether each response was correct or incorrect. These data were analyzed using a binomial general linear model which examined whether the probability of producing a successful response was affected by the time the participant had to prepare their response, the session in which the data was collected (Initial or Final), and the group to which the participant was assigned (Minimal Repeats or Frequent Repeats), with the formulation Correct ~ PreparationTime*Session*Group+(1|Subject). Significant interactions were analyzed using further binomial general linear models comparing the performance across sessions and between groups. To visualize the data, responses were binned over a 100ms window, and accuracy was determined as the proportion of correct responses within the 100ms window (Figure 1f) (Haith et al. 2016; Hardwick et al. 2019). We generated 95% confidence intervals using a resampling with replacement bootstrapping procedure with 10000 resamples. When a resample was drawn the same pool of participants was used to generate a pre and post training group average, effectively providing a repeated-measures assessment (Hardwick et al. 2017; Hardwick and Celnik 2014).

We also assessed participant’s general ability to perform the forced response task using two measures. We first examined the ability of participants to time their responses to occur with the fourth tone. Response asynchrony was assessed by subtracting the time at which participants responded from the time at which the fourth tone occurred. Response asynchrony was analyzed using a general linear model with the formulation Asynchrony ~ Session*Group+(1|Subject). Secondly, we measured the percentage of forced response trials in which participants failed to respond (i.e. no response was recorded more than 300ms after the final tone). The percentage of trials in which participants failed to respond was analyzed using a mixed model ANOVA with a between-participants factor of group (Minimal Repeats or Frequent Repeats) and a within-participants factor of session (Initial or Final).

### Rapid Response Trials

During the rapid response trials, performance was measured using reaction times and errors. Reaction time for each trial was defined as the time between stimulus onset and the time that a participant made their first response. These data were analyzed using a linear mixed model with the formulation: Reaction Time ~ Group*Session+(1|Subject). As we conducted a separate analysis of errors during rapid response trials, the analysis of reaction times examined only trials in which the first response the participant made was correct. The analysis of participant errors took the number of errors made in each block of the experiment and examined whether the probability of producing an error was affected by the group or session in which the block was completed, with the formulation: Errors ~ Group*Session+(1|Subject).

We also conducted analyses to determine whether repeating the same trial type on consecutive trials affected performance during training. Separate analyses were conducted for reaction times and errors. We first analyzed reaction times using a model with the formulation Reaction Time ~ Group*Session*Repeat+(1|Subject). For the analysis of errors, to control for the difference in the total number of repeat and non-repeat trials that each group experienced, these data were analyzed as percentages of the total number of repeat or non-repeat trials within each block. A mixed model ANOVA examined the between-participants factor of group (Frequent Repeats Group, Minimal Repeats Group), and the within-participants factors of trial type (Repeat, Non-Repeat) and session (1-5). To account for the very small number of repeat trials for the Minimal Repeats Group, we took the median average of the data from the blocks within each session, which provided an estimate of average performance that is more robust to outliers.

Changes in individual participant performance were illustrated by determining the change in the average preparation time required to produce a correct response. A previously established model (Haith et al. 2016; Hardwick et al. 2019; see Supplementary Materials) was used to determine the average preparation time that participants required to produce a correct response for the initial assessment, and the final assessment. The difference between the average time required to produce a correct response was used to measure the change in participant performance between the initial and final sessions.

## RESULTS

### Criterion Test Trials

During the criterion training block, participants learned the mapping of the symbols to the different keys. The groups did not differ significantly by the number of trials they needed to reach the criterion (t_29_=1.19, p=0.24) (see Figure 2a), nor by their reaction times when completing these trials (χ^2^=0.47, p=0.49) (see Figure 2b). Consequently, there was no significant difference between the groups at baseline when they learned the stimulus-response associations.

**Figure 2.**
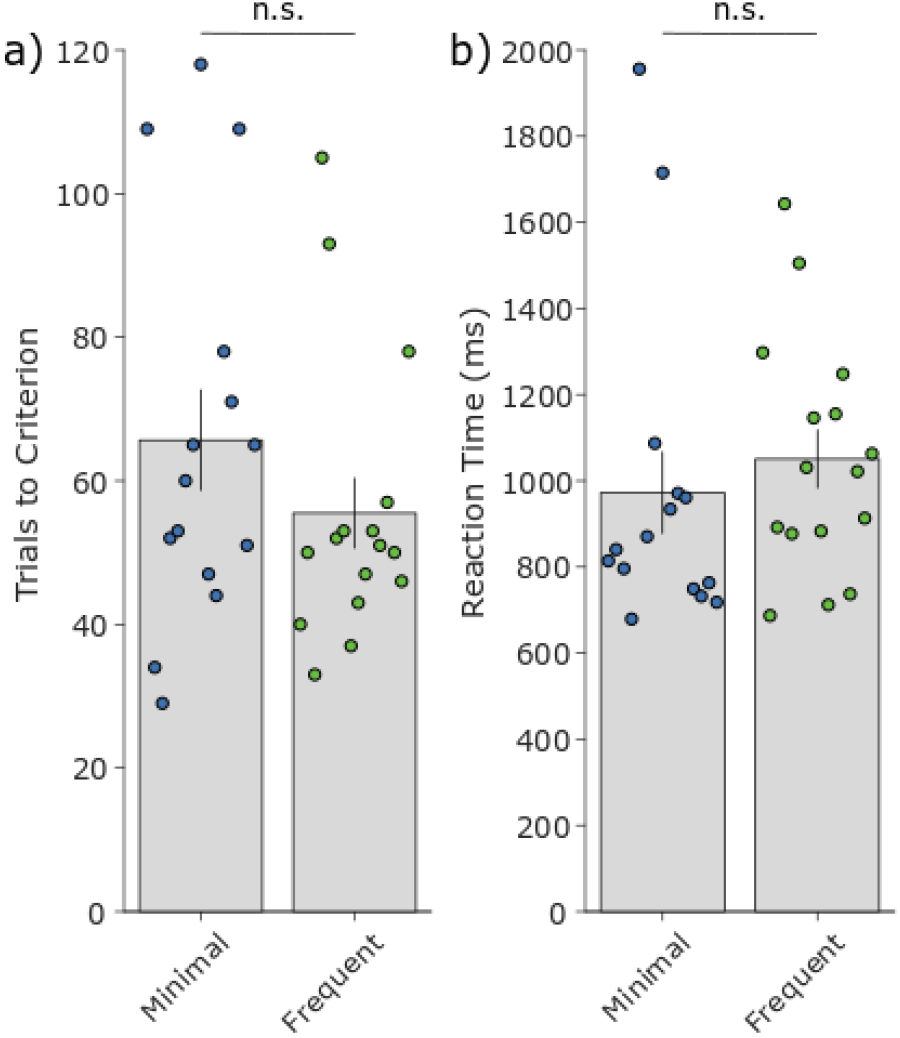
<u>Criterion measures</u>. a) The number of trials participants needed to reach the criterion. b) The average reaction times on the trials during the criterion training block. Grey bars represent group averages. Error bars represent the standard error of the mean. Dots represent data from individual participants. In all panels Minimal Repeats Group N=15, Frequent Repeats Group N=16.

### Forced Response Trials

Speed-accuracy trade-off curves were generated using data from the forced response trials collected on the first and last day of the experiment. A general linear model identified a significant three-way interaction between preparation time, session, and group (χ^2^=6.16, p=0.01). Participants in both groups required significantly less time to produce correct responses following training (general linear model comparing initial vs final performance for the Minimal Repeats Group, χ^2^=56.04, p=7.116e-14, and the Frequent Repeats Group, χ^2^=23.62, p=1.175e-06). While there was no significant difference between the groups at the initial assessment (χ^2^=0.76, p=0.38), critically, following their differing training interventions, the Minimal Repeats Group required significantly less time to select correct responses than the Frequent Repeats Group (χ^2^=5.99, p=0.01) at the final assessment. Model fits to individual data found that participants in the Minimal Repeats Group reduced the average time needed to produce accurate responses by 74±10ms (mean±SEM), while participants in the Frequent Repeats Group reduced this time by only 41±10ms (Figure 3e-g). To control for outliers (i.e. one participant in the Frequent Repeats Group that showed a decline in performance from the initial to the final session rather than an improvement), we conducted a sensitivity analysis. We found that the overall pattern of our results did not change if this participant was excluded from our analyses (the empirical result of the final assessment difference between the forced response data for the Frequent Repeats and Minimal Repeats Groups remained consistent: χ^2^ = 6.68, p = 0.0097). Thus, our empirical result stands and is robust to this potential outlier.

**Figure 3.**
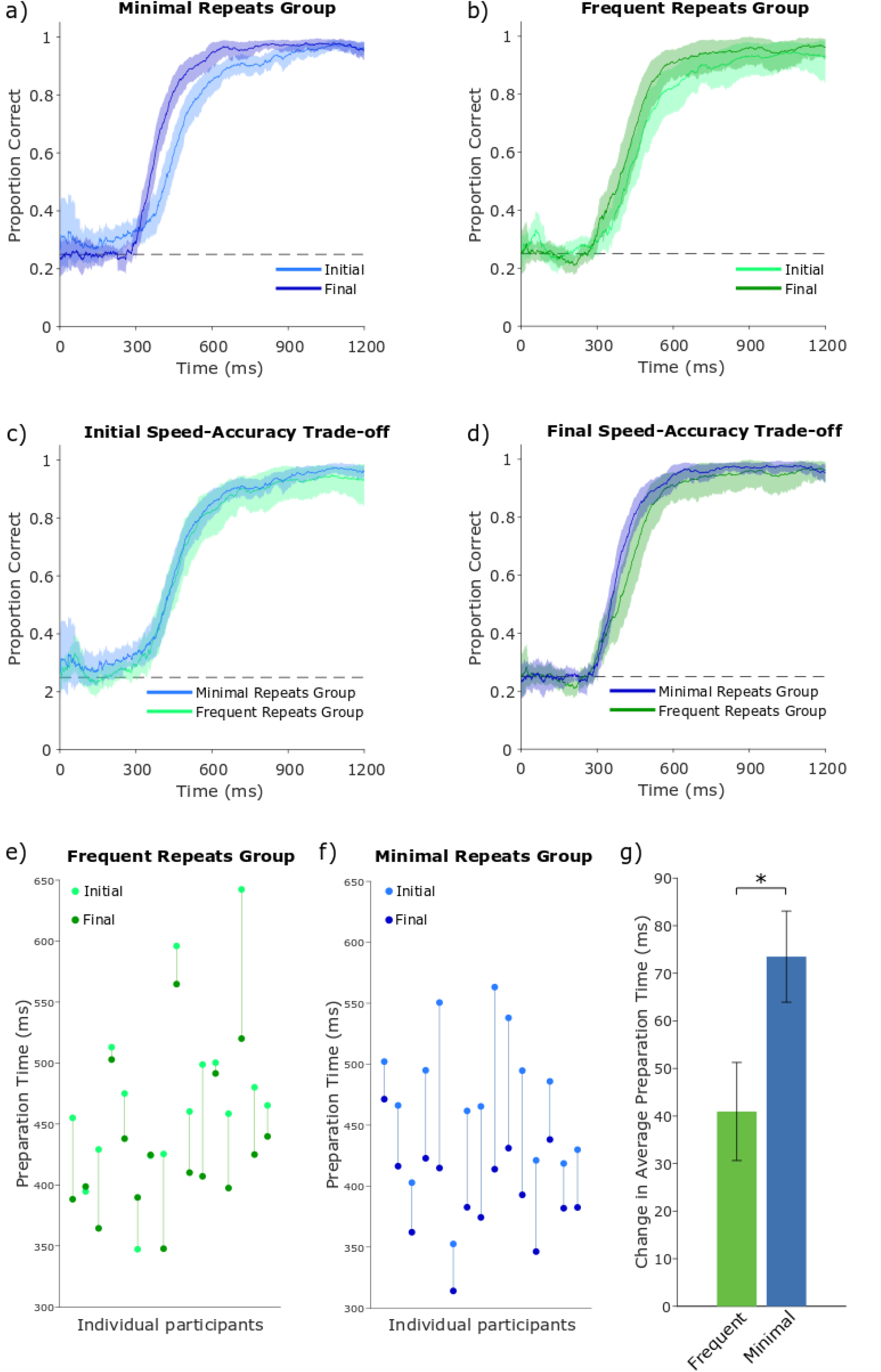
<u>Speed-accuracy trade-offs</u>. a) Initial versus final comparison of the speed-accuracy trade-off for the Minimal Repeats Group. b) Initial versus final comparison of the speed-accuracy trade-off for the Frequent Repeats Group. c) Comparison of the initial speed-accuracy trade-offs between groups. d) Comparison of the final speed-accuracy trade-offs between groups. In panels a-d shaded areas represent bootstrapped 95% confidence intervals. Dashed line indicates chance (1/4) level of performance. e,f) Summary data from model fits to individual participant performance (see also Supplementary Materials). Each line represents a single participant; the light circle represents the average time that a participant required to prepare a correct response in the initial session, and the dark circle represents this time in the final assessment. Line length therefore presents the change in average preparation time for that participant from the initial assessment to the final assessment. g) Group summary presenting the mean change in the average preparation time (i.e. average of line lengths in e-f). In all panels Frequent Repeats Group N=16, Minimal Repeats Group N=15.

**Figure 4.**
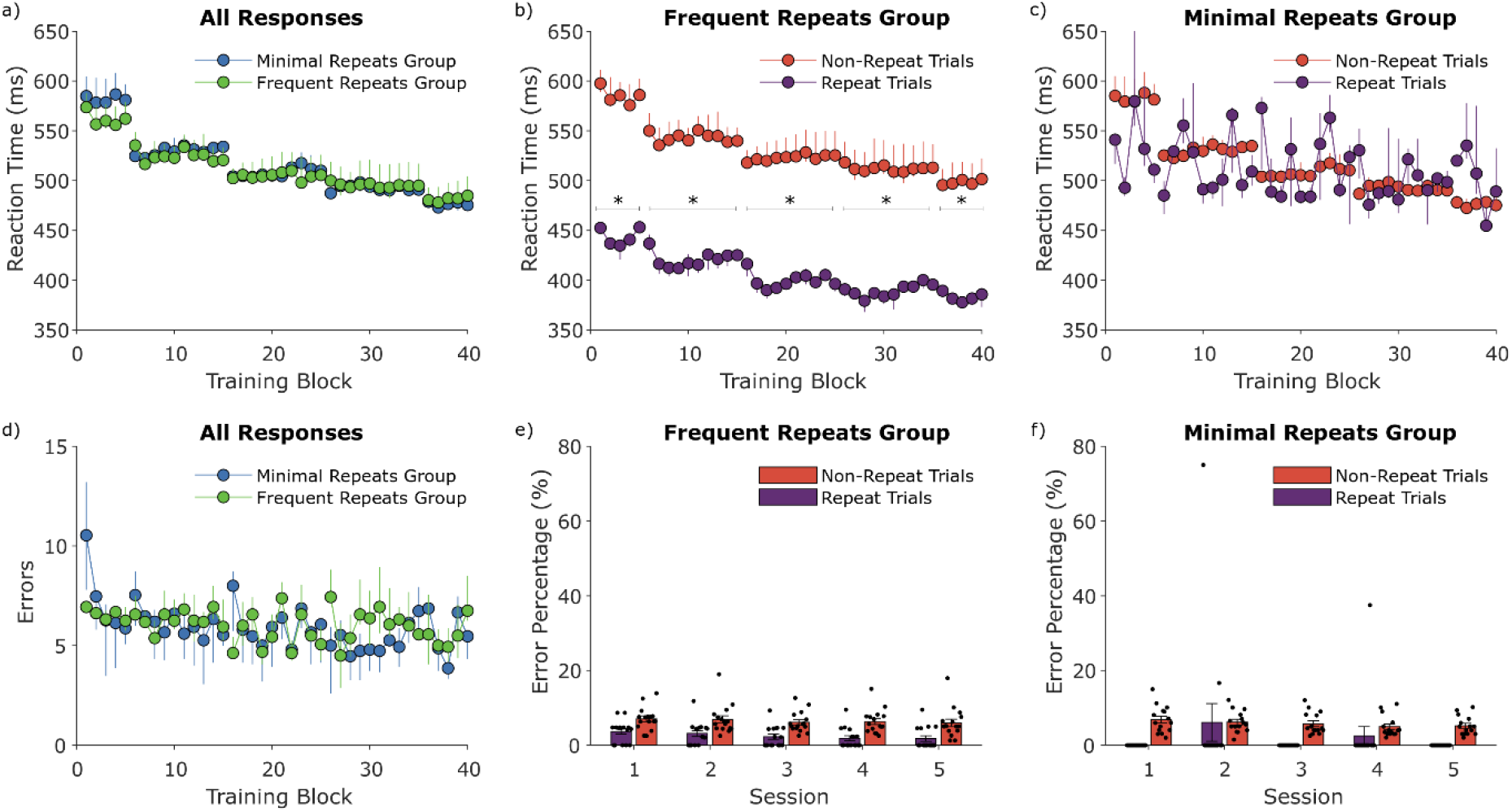
<u>Training Measures</u>. a) Plot of the reaction times during rapid response trials for both the Minimal Repeats (blue) and the Frequent Repeats (green) Group. Each point represents the average reaction time for correct trials within a 100-trial block. b) Plot of the reaction times during rapid response trials for both non-repeat (red) and repeat (purple) trials of the Frequent Repeats Group. c) Plot of the reaction times during rapid response trials for both non-repeat (red) and repeat (purple) trials of the Minimal Repeats Group. d) Plot of the number of first attempt errors during rapid response trials for both the Minimal Repeats (blue) and the Frequent Repeats (green) Group. e) Plot of the percentage of errors during rapid response trials for both non-repeat (red) and repeat (purple) trials of the Frequent Repeats Group. f) Plot of the percentage of errors during rapid response trials for both non-repeat (red) and repeat (purple) trials of the Minimal Repeats Group. In figures a, b, c, and d, error bars represent bootstrapped 95% confidence intervals. In figures e and f (bar charts), error bars represent the standard error of the mean. In all panels Minimal Repeats Group N=15, Frequent Repeats Group N=16.

Further analyses examined the ability of participants to perform the general requirements of the forced response task. Participants typically produced their response just prior to the fourth tone, and their timing was more precise in the final session (significant effect of session, χ^2^=187.25, p<2e-16; initial session: 14±5ms, final session: 7±4ms, mean±SEM). There was no significant difference in response asynchrony between groups (χ^2^=0.00, p=0.96), nor was there a significant group by session interaction (χ^2^=0.18, p=0.67). Analysis of the percentage of forced response trials in which participants failed to respond revealed that participants failed to respond on fewer trials in the final session (mixed model ANOVA, main effect of session, F=7.02, p<0.05; initial session: 0.00226±0.00326% of trials (mean±SEM), final session: 0.00083±0.00005% of trials, with no significant difference between groups; Group*Session interaction, F=1.92, p=0.18).

### Rapid Response Trials

A linear mixed model examining participant’s reaction times during the training period revealed a significant effect of session (χ^2^=6691.73, p<2e-16), showing that participants improved their performance with practice. The linear mixed model also revealed a significant group by session interaction (χ^2^=176.88, p<2e-16); however, further simple main effects analysis comparing the performance of the two groups on each of the five days of the experiment revealed no significant differences (Session 1: χ^2^=1.70, p=0.19, Session 2: χ^2^=1.10, p=0.29, Session 3: χ^2^=0.91, p=0.34, Session 4: χ^2^=0.07, p=0.79, Session 5: χ^2^=0.01, p=0.92).

The second variable of interest for the training blocks was the number of errors. A linear mixed model identified a significant effect of session (χ^2^=25.24, p=4.504e-05), with participants reducing the number of errors they made with training. While there was a trend toward significance for the interaction between group and session (χ^2^=10.57, p=0.03), simple main effects analysis examining performance of the two groups on each session of the experiment revealed no significant differences between the groups for each session (Session 1: χ^2^=0.39, p=0.53, Session 2: χ^2^=0.03, p=0.87, Session 3: χ^2^=0.13, p=0.72, Session 4: χ^2^=0.58, p=0.45, Session 5, χ^2^=0, p=1).

A secondary analysis examined differences between repeat and non-repeat trials on performance. Overall, reaction times were significantly faster when participants responded to repeat trials compared to non-repeat trials (significant effect of Repetition, χ^2^=9131.81, p<2.2e-16). As this analysis revealed a significant three way interaction between Group, Session, and Repetition (χ^2^=13.86, p<0.01), we conducted further analyses for each group. Participants in the Frequent Repeats Group were faster for repeat trials compared to non-repeat trials within each session (Session 1: χ^2^=1006, p<2.2e-16, Session 2: χ^2^=2786.60, p<2.2e-16, Session 3: χ^2^= 3070.40, p<2.2e-16, Session 4: χ^2^=2824.10, p<2.2e-16, Session 5: χ^2^=1628.30, p<2.2e-16). Initially, participants in the Minimal Repeats Group were also significantly faster when responding to repeat trials compared to non-repeat trials (Session 1: χ^2^=15.05, p<0.001, Session 2: χ^2^=5.43, p<0.02), but this effect was not present in later sessions (Session 3: χ^2^=0.45, p=0.50, Session 4: χ^2^=0.003, p=0.96), and showed a trend for reversal in the final session (Session 5: χ^2^=3.42, p=0.06).

Analysis of the percentage of errors made in repeat vs non-repeat trials revealed a significant main effect of trial type (F_1,29_=72.73, p<0.001), indicating that participants were significantly less likely to make errors in repeat trials compared to non-repeat trials. There was also a trend for a reduction in errors with practice (effect of Session, F_4,116_=2.07, p=0.089). No significant interaction effects were present, indicating that these effects did not differ between groups (all F<1.3, p>0.25).

## DISCUSSION

Here we examined the impact of repeating consecutive trials during practice on learning using an Arbitrary Visuomotor Association Task. We found that after practice, the Minimal Repeats Group, who frequently experienced different stimuli on consecutive trials, learned more than the Frequent Repeats Group, who often experienced repeating stimuli on consecutive trials.

A speed-accuracy trade-off measure was used to assess skill acquisition. While both groups needed less time to obtain a given level of accuracy after five days of practice, the Minimal Repeats Group improved to a greater extent than the Frequent Repeats Group. This is in line with previous studies investigating the impact of blocked versus random practice on performance (Goode and Magill 1986; Magill and Hall 1990; Pauwels et al. 2014; Smith and Davies 1995). One possible explanation for this effect was that the Minimal Repeats Group trained on a harder task than the Frequent Repeats Group. Evidence from the training measures of the Frequent Repeats Group shows that both the reaction times and error rates were lower for repeat than non-repeat trials. This is consistent with previous research indicating that repeating the same response facilitates performance (Bertelson 1961; Maljkovic and Nakayama 1994, 1996; Tanaka and Shimojo 1996). As described by the Challenge Point Framework, in order to learn more, you need to execute a harder task as this challenges you more (Guadagnoli and Lee 2004). In the framework of contextual interference, Pauwels et al. (2014) proposed that block-level contextual interference adds additional challenge to the learning process (i.e. increased cognitive effort and processing) which has beneficial effects on learning. Hence, as contextual interference at the individual trial level makes the task harder, it would have similar beneficial effects on learning in the present experiment. So, whereas previously it has been shown that repetition leads to short-term (trial-to-trial) benefits, we now highlight that trial-to-trial repetition may not be as efficient as trial-to-trial switching in inducing longer term learning benefits.

While both groups differed with regards to the speed-accuracy trade-offs from the forced response trials, the main analysis of their training data did not reveal differences between their reaction times or error rates. There are, however, multiple explanations for this apparently paradoxical result. First, it is important to note that both groups trained on different versions of the task. As discussed above, the training task was easier for the Frequent Repeats Group, as was confirmed by our secondary analyses of the effects of repetition on performance during the rapid response training trials. As such, a direct comparison between the performance for each group during training may not reveal an actual performance difference between them. A further issue is that it can be difficult to interpret the data from a task where speed and accuracy are considered separately due to the ‘performance confound’. As speed and accuracy are both inherently linked to the overall performance of a skill (Fitts 1954), this allows some variability between participants. While participants were instructed to respond as quickly and as accurately as possible to the symbols, this instruction can be interpreted differently by each subject; some might focus more on speed at the cost of accuracy, while others might favor accuracy at the cost of speed (note that the variability of participant responses in the criterion assessment condition illustrates this effect). This illustrates the point that it is difficult to assess skill when interpreting data where both speed and accuracy can vary freely (Hardwick et al. 2017; Reis et al. 2009; Shmuelof et al. 2012). Hence, Reis et al. (2009) suggested that the performance of a skill should be quantified using a speed-accuracy trade-off. Therefore, in the present study, the forced response task was used to provide a clearer assessment of overall performance; controlling the time participants had to prepare their responses (by varying the onset of the visual stimulus), allowed us to assess accuracy without the influence of confounding differences in response speed.

There has been much research investigating the effects of repetition on learning through contextual interference (for reviews see Brady 1998; Magill and Hall 1990; Merbah and Meulemans 2011). By comparison, to the best of our knowledge, studies investigating trial-to-trial effects of repetition on *overall* learning have not been subject to much research (i.e. only one study using a between-subjects design; Kaipa et al. 2016, which involved only two response alternatives). This seems surprising as there has been great interest in developing approaches to enhance learning. Notably, previous work has tried to improve learning rates using relatively complex and more direct neuromodulatory approaches including non-invasive brain stimulation (Galea et al. 2011; Hardwick and Celnik 2014; Reis et al. 2009; but see also López-Alonso et al. 2015, 2018; Vallence et al. 2013) and pharmacological drug interventions (Pugliese 1973). Behavioral manipulations such as the approach highlighted in the present study provide simple and cost effective alternatives to these direct neuromodulatory approaches, and are also free from their inherent increased risk to participants (Reis et al. 2008; Rossi et al. 2009). Furthermore, there is potential to combine both direct neuromodulatory and behavioral interventions to further enhance the learning process, which could result in larger performance benefits.

Our results showed that practicing with few repeating consecutive trials led to a greater skill improvement. Participants practiced the task for several thousand trials, and were able to improve their ability to produce short-latency responses, consistent with previous work indicating that relatively high volumes of training allow participants to produce automatic, habitual responses (Hardwick et al. 2019). As skill acquisition requires extensive practice (Ericsson et al. 1993), developing approaches to enhance the rate of learning is of interest across a diverse range of fields such as sports, playing musical instruments, and learning surgical skills. Enhancing the rate of learning could also be applied to rehabilitation, and may specifically be beneficial during the early stages of recovery following neurological insult. For example, most recovery from stroke is made within the “acute phase” within the first three months after the infarct (Xu et al. 2017). During this critical time period, the brain naturally reorganizes itself by means of neuroplastic mechanisms, which leads to a period of “spontaneous biological recovery” (Zeiler and Krakauer 2013). It has therefore been argued that training during this critical period would result in an improved overall recovery due to the spontaneously occurring neuroplastic processes interacting with physical practice (Zeiler and Krakauer 2013). Taken together, this underlines the importance of developing approaches to improve the rate of learning.

Previous work indicates that motor skill learning can occur at the levels of action selection (changes in ability to select an appropriate action) and action execution (changes in the quality of the performed action) (Chen et al. 2018; Diedrichsen and Kornysheva 2015). The Arbitrary Visuomotor Association Task used in the present study primarily focuses on action selection, assessing the ability to quickly and accurately choose an appropriate response. We believe that our training manipulation led to a specific enhancement in this response selection ability. Prior evidence indicates that it is more taxing to select different actions from trial-to-trial than it is to repeat the same action (Fecteau 2007; Gupta and Cohen 2002; Maljkovic and Nakayama 1994, 1996; Tanaka and Shimojo 1996). As such, we propose that even though the two groups in the present study completed the same number of training trials, the mechanisms involved in response selection were more frequently engaged to their full extent by the task presented in the Minimal Repeats condition compared to the Frequent Repeats condition. It remains to be seen what effect, if any, manipulating the number of trial-to-trial repeats would have on the action execution component of motor skill learning.

Research investigating the neural correlates of action priming suggests that the repetition of a stimulus leads to suppression of neuronal responses in the brain (i.e. reduced firing rate of neurons) (Desimone 1996; Henson and Rugg 2003; Wiggs and Martin 1998). Research in human neuroimaging indicates an analogous reduction in the brain’s haemodynamic response when participants repeat the same action across consecutive trials (Hamilton and Grafton 2009), which has been attributed to a similar decrease in neuronal firing (Bunzeck and Thiel 2016; Grill-Spector et al. 2006). Given that neuronal co-activity has been proposed as the primary mechanism of neuroplasticity (Hebb 1949), this effect has important implications for learning. While repeating the same action facilitates short-term, trial-to-trial performance, we speculate that the reduction in neuronal firing that occurs with repetition may, in the longer term, reduce the rate at which connections in the brain are formed and strengthened, consequently reducing the overall rate of learning. Such an effect could readily explain the present findings, and is consistent with findings in previous research. For example, Chalavi et al. (2018) found that random practice led to decreased GABA levels, while blocked practice was associated with increased levels of GABA. As disinhibition via GABA reduction has been suggested to be a primary mechanism whereby the brain forms new motor memories (Stagg et al. 2011), this suggests that repeating the same action as occurs during blocked practice indeed has a net-negative effect on neuroplasticity. As such, we believe this offers a plausible explanation for our results, though further research using measures of brain activity during learning and/or measures of brain connectivity following learning is required to further complement this proposal.

### Strengths and Limitations

The present study demonstrates an important point for learning experiments that present trials in a random order; repeating the same action is easier than switching between different actions, and reduces the rate at which participants learn. Notably, for both groups, the frequency of repeat trials was lower than may be expected from pure random ordering of trials, which on average would occur on 24 trials in a 100 trial block. Here, our Frequent Repeats Group were presented with repeat trials in a pseudorandom order to ensure that participants would be exposed to a similar number of repeat trials for each stimulus. Critically, even though this led to fewer repeat trials per block than may occur in a purely random trial order, our results still show a significant detrimental effect of trial-to-trial repetition on learning. As such, the frequency of repeat trials for the Frequent Repeats Group, even if it is lower than might be anticipated through fully random trial ordering, serves the purpose of illustrating that trial-to-trial repetition is not favorable with regards to learning and performance.

Participants in the present study trained over multiple days, completing the first and final session of the study while being supervised by an experimenter in the laboratory. The intervening sessions were conducted alone at their home. As such, it could be argued that practicing the task in these different contexts could have had an impact on participants’ performance (i.e. the presence of an experimenter has previously been shown to affect participants’ performance; for a review see McCambridge et al. 2014). However, as this aspect of the procedure did not differ between groups, it appears unlikely that it could account for our between-group differences. As such, allowing participants to train at home provided a pragmatic solution to conduct an experiment across multiple days. In the future, the paradigm used here could be extended by increasing the number of days over which participants trained, and/or including assessment of retention of learning after a period without training, which has previously been shown to be greater in groups that experienced contextual interference (Battig 1979; Cross et al. 2007; Goode and Magill 1986; Pauwels et al. 2014; Smith and Davies 1995).

While the Frequent Repeats Group consistently showed a beneficial effect of repetition on response times during training, the pattern of results for the Minimal Repeats Group differed. While the Minimal Repeats Group also had faster response times to repeating stimuli in the first two sessions of the experiment, this effect was not present in sessions three and four, and there was a non-significant trend for participants to be faster for non-repeat trials by session 5. This effect may indicate that, because participants in the Minimal Repeats Group rarely experienced repeat trials, they did not gain beneficial effects from them in later blocks due to a lack of exposure. However, given the extremely small sample size of repeat trials in the Minimal Repeats Group, we treat this result with caution, and suggest further experimentation is required before a firm conclusion can be drawn on this point.

Participants in the present study were not informed that two separate groups were completing the experiment, and were not told that we were examining the effects of the frequency of trial-to-trial repetition on performance. However, we did not assess whether participants became aware of the experimental manipulation during the study. While we believe this to be unlikely due to the relatively subtle nature of the intervention, we do not know how (if at all) explicit awareness of the manipulation would affect performance. Future studies may wish to perform such manipulation checks to examine this effect.

Finally, the present experiment used a relatively ‘artificial’ arbitrary visuomotor association paradigm. Future research is required to determine whether the effects presented here would also occur in more ecologically valid paradigms such as training processes in real-life settings such as sports, music, and rehabilitation. However, as arbitrary associations underlie everyday skills such as language, typing, and driving, and as previous research has shown beneficial effects of contextual interference not only in laboratory paradigms (Battig 1979; Cross et al. 2007; Pauwels et al. 2014), but also in real-world skills such as as kayaking (Smith and Davies 1995) and badminton (Goode and Magill 1986), we speculate that our present findings would generalize across a wide range of settings.

## CONCLUSIONS

Here we investigated the effect of switching between trials versus repeating of consecutive trials during practice on learning. We found that, following training, a group who trained with minimal trial-to-trial repetition showed greater improvement than a group who frequently experienced trial-to-trial repetition. As such, our research indicates that while repetition benefits short-term performance, minimizing trial-to-trial repetition leads to greater long-term skill acquisition. This simple but effective manipulation of practice could be readily applied in the contexts of skill learning in fields such as sport, music, and rehabilitation.

## AUTHOR CONTRIBUTIONS

RMH designed the experiment. LWEV collected the data. LWEV, in collaboration with RMH, analyzed the data. LWEV drafted the manuscript. LWEV, SPS, and RMH reviewed and edited the manuscript.

